# Generation and Functional Characterization of PLAP CAR-T Cells against Cervical Cancer Cells

**DOI:** 10.1101/2022.06.12.495798

**Authors:** Vahid Yekehfallah, Saghar Pahlavanneshan, Ali Sayadmanesh, Zahra Momtahan, Bin Ma, Mohsen Basiri

## Abstract

Chimeric antigen receptor (CAR) T cell therapy is one of the cancer treatment modalities that has recently shown promising results in treating hematopoietic malignancies. However, one of the obstacles that need to be addressed in solid tumors is the on-target/off-tumor cytotoxicity due to the lack of specific tumor antigens with low expression in healthy cells. Placental alkaline phosphatase (PLAP) is a shared placenta/tumor-associated antigen (TAA) that is expressed in ovarian, cervical, colorectal, and prostate cancers and is negligible in normal cells. In this study, we constructed second-generation CAR T cells with a humanized scFv against PLAP antigen, then evaluated the characteristics of PLAP CAR T cells in terms of tonic signaling and differentiation in comparison with ΔPLAP CAR T cells and CD19 CAR T cells. In addition, by coculturing PLAP CAR T cells with HeLa cells, we analyzed the tumor-killing function and secretion of anti-tumor molecules. Results showed that PLAP CAR T cells not only could eliminate the cancer cells but also increase their proliferation in vitro. We also observed increased secretion of IL-2, granzyme A, and IFN-γ by PLAP CAR T cells upon exposure to the target cells. In conclusion, PLAP CAR T cells are potential candidates for further investigation in cervical cancer and potentially other solid tumors.

## 1. Introduction

Immune cells can detect cancer cells and eliminate them by patrolling the individual’s body, but in case of a cancer outbreak, immune cells’ anti-tumor effect does not have enough capacity to recognize and overcome cancer cells [1]. Hence, using modalities to enhance the potential of immune cells is imperative. For this goal, one of the methods that can be used is integrating the variable regions of antibodies, and signaling domains from T cells, to make a chimera, known as chimeric antigen receptor (CAR) T cells [2].

CAR T cells have demonstrated satisfactory results in hematopoietic malignancies [3, 4]. Especially, in a recent study by Melenhorst *et* al., they found that CD19-specific CAR T cells not only have anti-tumor capabilities against chronic lymphocytic leukemia but also show persistency and signaling in CD19 CAR T cells that can act as living drugs in 2 patients with leukemia, a decade after initial treatment [5]. Nevertheless, CAR T cells need to overcome some obstacles in solid tumors in order to perform their anti-tumor effects [6]. One of these obstacles is on-target/off-tumor toxicity; in this situation, CAR T cells detect the shared targeted antigens on healthy cells instead of cancer cells, so discovering the tumor antigen specifically expressed on cancer cells can address this side effect [7].

Recent investigations [8, 9] show that placental alkaline phosphatase (PLAP) can be a potential antigen for targeted cancer immunotherapy. In fact, PLAP or alkaline phosphatase, placental type (ALPP), is a glycosylated membrane-bound enzyme that is normally expressed in the placenta and testis, and will be increased in malignancies including ovarian [8], prostate [9], cervical [10], colorectal, and testicular seminoma [11]. The other characteristics of PLAP which make it a potential target for cancer immunotherapy include the accessibility of PLAP, low expression in healthy cells, and high activity of PLAP in late-stage cancers, resulting in tumor progression [12]. Interestingly, Fushimi *et* al., have proven that intervening in PLAP expression can prohibit calcification of breast cancer cells[13].

Owing to the high expression of PLAP in cervical cancers [14, 15] and the importance of cervical cancer as the second most common malignant disease in women [16, 17], we hypothesized that PLAP could be a potential target for CAR T cells [18]. Hence, in this experiment, we aim to design and construct second-generation CAR T cells against PLAP^+^ cells in cervical cancer cells, including HeLa cells. In order to assess the anti-tumor cytotoxicity of PLAP CAR T cells, we compared their characteristics with CD19 CAR T cells in terms of tonic signaling and differentiation markers. Furthermore, we demonstrated that PLAP CAR T cells could be prospective CAR T cells in solid tumors.

## 2. Materials and Methods

### 2.1 Cell Lines

Plat-A, HeLa, and HEK293 cell lines were obtained from Royan Institute Cell Bank and cultured in Dulbecco’s Modified Eagle Medium (DMEM, GE Healthcare Life Sciences, Pittsburgh, PA, USA) supplemented with 10% heat-inactivated fetal bovine serum (FBS; Thermo Fisher Scientific, Gaithersburg, MD, USA)1% penicillin/streptomycin (Invitrogen, Eugene, OR, USA) and 2 mM L-GlutaMAX (Thermo Fisher Scientific). All cell lines were grown in 85% humidity with 5% carbon dioxide (CO2) at 37 °C.

### 2.2 Generation of Retroviral Constructs and Retroviral Vectors

PLAP CAR was designed based on published sequences of PLAP-specific single-chain variable fragment (scFv) [19], human CD8a signal peptide, cMyc-tag, human CD28, and CD3ζ proteins. The sequence was optimized for expression in human cells and then ordered for synthesis. First, the synthetic sequence was cloned into the SFG vector using *Nco*I and *Sph*I (Thermo Fisher Scientific) restriction enzymes. Correct clones were confirmed by sequencing the inserted CAR sequence. Next, to produce the ΔPLAP CAR construct, the signaling portion of CD28 and CD3ζ endodomain were excised using the *Xho*I restriction enzyme, and the plasmid was recirculated by T4 DNA ligase (Thermo Fisher Scientific). The CAR constructs were transfected to Plat-A packaging cell line via Lipofectamine 3000 reagent (Thermo Fisher Scientific) and retroviral supernatant was collected at 48 and 72 hours post-transfection, then filtered (using a 0.45-mm filter) and stored at –80 °C.

### 2.3 Generation of CAR T Cells

Written informed consent was collected before isolating peripheral blood mononuclear cells (PBMCs) by ficoll density centrifugation from healthy donor volunteers. The protocols were approved under the Research Ethics Committee at Royan Institute. 1 × 10^6^ PBMCs were seeded in each well of a non-tissue culture-treated 24-well plate that had been pre-coated with OKT3 (1 mg/ml) (Ortho Biotech, Inc., Bridgewater, NJ, USA) and CD28 (1 mg/ml) (Becton Dickinson & Co., Mountain View, CA, USA). On day one, cells were grown in complete media [RPMI-1640 containing 45% Clicks medium (Irvine Scientific, Inc., Santa Ana, CA, USA), 10% FBS, and 2 mM L-GlutaMAX], which contained recombinant human IL2 (100 U/mL, Royan Biotech, Tehran, Iran). On day 3, after OKT3/CD28 T blast generation, 1 mL of retroviral supernatant was added to a non-tissue culture-treated 24-well plate pre-coated with Retronectin (FN CH-296; Takara Shuzo, Otsu, Japan) and centrifuged for 90 min at 2000×g. OKT3/CD28 activated T cells (0.2 × 10^6^/mL) were resuspended in complete media supplemented with IL2 (100 U/mL) and then added to the wells and centrifuged at 400×g for 5 min. Transduction efficiency was measured 3 days post-transduction by flow cytometry.

### 2.4 Flow Cytometry Analysis

For analyzing surface markers with flow cytometry, cells were collected, washed, and stained with antibodies (Table 1) at 4°C for 30 min in the dark. For intracellular staining of phospho-CD3ζ, T cells were fixed with 1.5% formaldehyde solution (F1635, Sigma-Aldrich, St. Louis, MO), washed, permeabilized with pre-chilled 100% methanol (Fisher Scientific, Pittsburgh, PA) on ice for 15 min, and then washed three times, followed by staining with CD247(pY142)-specific antibody (Table1) for 60 minutes at room temperature, in the dark. All samples were acquired on either BD FACSCalibur or BD FACSCanto flow cytometers (BD Biosciences, USA), and the data were analyzed with FlowJo software (Tree Star Inc., Ashland, USA).

**Table 1.** Antibodies used for flow cytometry assays.

| Target Protein | Clone | Fluorophore | Company | Cat. No. |
| --- | --- | --- | --- | --- |
| cMYC | 9E10 | FITC | Sigma-Aldrich | F2047 |
| CD3 | UCHT1 | APC-Alexa Fluor 750 | Beckman Coulter | A66329 |
| CD4 | 13B82 | Krome Orange | Beckman Coulter | A96417 |
| CD8 | SK1 | Pacific Blue | BioLegend | 344718 |
| CD45RA | 5H9 | APC | BD Biosciences | 561210 |
| CCR7 | 150503 | Alexa Flour 700 | BD Biosciences | 561143 |
| CD27 | O323 | PE Cy7 | BD Biosciences | 567290 |
| CD28 | CD28.2 | PE Cy5 | BD Biosciences | 561791 |
| CD247(pY142) | K25-407.69 | Alexa Fluor 647 | BD Biosciences | 558489 |
| CD69 | TP1.55.3 | ECD | Beckman Coulter | 6607110 |
| CD25 | M-A251 | PE Cy5 | BD Biosciences | 555433 |

### 2.5 Cytotoxicity assay

Luciferase-expressing HeLa or HEK293 cells were seeded into a U-bottomed 96-well plate plus D-Luciferin (75 μg/ml; Sigma-Aldrich) for cytotoxicity assessment. After that, CAR-expressing T cells were added to the mix and cells were grown and incubated at 37 °C in 5% CO2 for the duration of the assay. The LUMIstar Omega microplate luminometer (BMG Labtech; Ortenberg, Germany) was used to monitor luminescence at every time point as relative light units (RLUs). In each assay, for baseline lysis, one group of target cells was cultured alone, and for maximum lysis, another group was cultured in 1% Triton X-100 (Sigma-Aldrich). Percentage of the specific lysis was calculated according to the following formula: % specific lysis = 100 × (spontaneous cell death RLU - sample RLU)/ (spontaneous death RLU – maximal killing RLU).

### 2.6 Activation assessment and cytokine measurement

To measure the amount of cytokine expression, 5×10^5^ transduced T cells were cocultured with HeLa target cells at a 1:1 ratio. After 24 hours, T cells and media were collected and separated by centrifugation. The cells were subjected to immunostaining with anti-CD25 and anti-CD69 and flow cytometry while the supernatants were stored at –20°C until assessed by ELISA for detection of secreted cytokines. ELISA assays were performed by Human Granzyme A DuoSet (DY2905-05, R & D Systems, MN, USA), Human IL-2 DuoSet (DY202, R & D Systems), and Human IFN-gamma DuoSet (DY285B, R & D Systems) as per the manufacturer’s instructions.

### 2.7 Coculture experiments

The coculture of 2×10^4^ T cells and 1×10^4^ GFP-FFLuc^+^ HeLa cells (2:1 ratio) was performed in a 12-well plate for 9 days in the presence of 2 mL complete media. For quantifying cells by flow cytometry, 20 μL of CountBright™ Absolute Counting Beads (C36950; Invitrogen) was added and 7-AAD was added for excluding the dead cells. The acquisition was halted at 2,500 beads.

### 2.8 Statistical analyses

Statistical significance between groups were determined using two-way ANOVA followed by Sidak’s multiple comparison test. P-values less than 0.0332 were considered statistically significant. Statistical analyses were performed using GraphPad Prism 8.0.2 (GraphPad Software Inc., La Jolla, CA, USA). All results are reported as mean ± SD.

## 3. Results

### 3.1. CAR construct and their expression on T cells

We designed a second-generation PLAP CAR containing a CD8α signal peptide, a human scFv against PLAP protein, a cMyc-tag, CD28 transmembrane cytosolic domains, and CD3ζ cytosolic domain. A non-signaling ΔPLAP CAR construct was used as negative control distinguished from PLAP CAR by the lack of CD28 and CD3ζ endodomains (Figure 1A). We checked the expression of CAR molecules on the surface of transduced cells 5-6 days after transduction using an anti-cMYC antibody that binds to the cMYC-tag of PLAP CAR and ΔPLAP CAR. Results showed that typically more than 80% of the transduced T cells expressed PLAP CAR and ΔPLAP CAR on their surfaces (Figure 1B).

**Figure 1.**
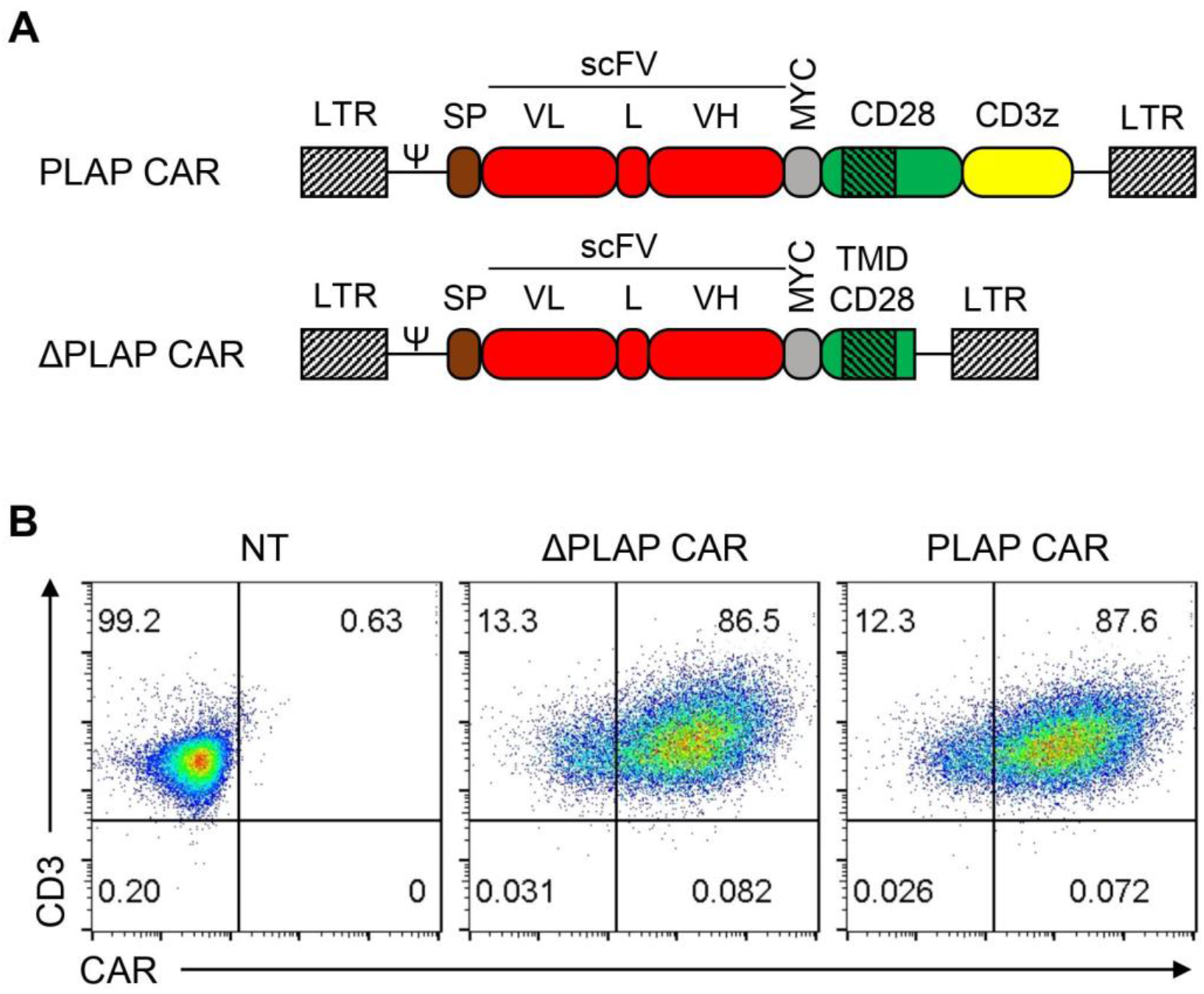
Design and expression of PLAP and ΔPLAP CAR constructs. (A) Scheme of PLAP CAR and ΔPLAP CAR. Both CARs contained a signal peptide, a codon-optimized synthetic gene encoding for humanized anti PLAP scFv, MYC tag and the transmembrane region of CD28. Intracellular signaling domains, including CD28 and CD3ζ, are only incorporated into PLAP CAR. (B) CAR expression was confirmed on primary T cells by flow cytometry.

### 3.2 Tonic signaling and immunophenotype of CAR T cells

Since antigen-independent tonic signaling can hamper CAR T cell anti-tumor activity, we first sought to determine any sign of tonic signaling or adverse effects on the immune phenotype of the expanded CAR T cells. As a positive control, we used a second-generation CD19 CAR containing the FMC.63 scFv, which is known to lack significant antigen-independent tonic signaling. We did not detect any antigen-independent growth in PLAP CAR T cells as compared with ΔPLAP and CD19 CAR T cells (Figure 2A). Furthermore, we used antibodies against the phosphorylated CD3ζ endodomain in the absence of PLAP antigen to directly assess antigen-independent signaling from the PLAP CAR molecule. According to our findings, ΔPLAP and PLAP CAR T cells like CD19 CAR T cells and non-transduced T cells did not phosphorylate in the CD3ζ domain; therefore, CAR T cells did not have tonic signaling (Figure 2B).

**Figure 2.**
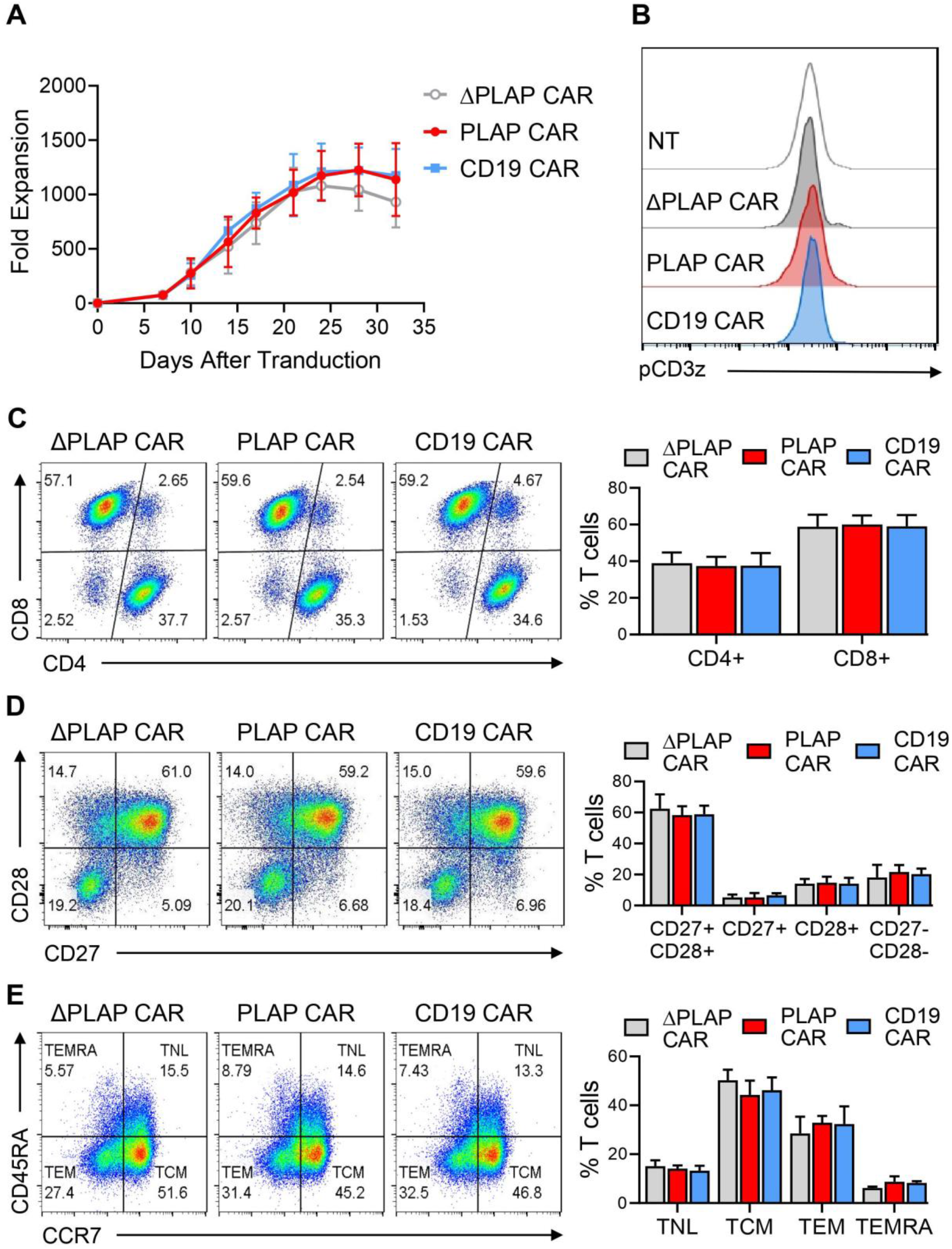
Tonic signaling and immunophenotype characterization of PLAP CAR T cells. (A) Lack of tonic signaling was confirmed by evaluating the growth of CAR T cells in the absence of antigen, and (B) no independent signaling in signaling domains. (C) The ratio of CD4:CD8 among CAR T cells was stable, which was assessedusing anti-CD4 and anti-CD8 antibodies by flow cytometry. (D) The population of CD27^+^CD28^+^ CAR T cells was more than reported presviously by anti-CD27 and anti-CD28 by flow cytometry. (E) TCM and TEM have more populations than TNL and TEMRA, performed by anti-CD45RA and CCR7. TNL is CD45RA^+^ CCR7^+^, TCM is CD45^−^CCR7^+^, TEM is CD45RA^−^ CCR7^−^, and TEMRA is CD45RA^+^ CCR7^−^. n=3.

We evaluated the CD4:CD8 ratio of CAR T cells by flow cytometry, observing no statistical significance between PLAP CAR T cells and the control groups (Figure2C). Moreover, we evaluated the expression of CD27 and CD28 costimulatory receptors on CAR T cells which are known to be correlated with T cell activity and less-differentiated naïve-like and memory phenotypes [20]. Results indicated that the majority (58.4±5.7%) of PLAP CAR T cells were CD27+CD28+ T cells. More importantly, there was no significant difference in the percentage of CD27 and CD28 expressing T cells between PLAP CAR T cells and the control groups (Figure 2D).

We also evaluated the expression of CD45RA and CCR7 markers on PLAP CAR, ΔPLAP CAR, and CD19 CAR T cells to survey the memory differentiation during *ex vivo* expansion. PLAP CAR T cells contained 14.1±1.3% naïve-like (TNL, CCR7^+^CD45RA^+^), 44.2±6.0% central memory (TCM, CCR7^+^CD45RA^−^), 32.9±2.8% effector memory (TEM, CCR7^−^CD45RA^−^), and 8.7±2.2% CD45RA-expressing effector memory T cells (TEMRA, CCR7^−^CD45RA^+^). We did not detect any significant difference in the percentage of these sub-populations among the experimental groups (Figure 2E).

Overall, these findings suggest that the PLAP CAR construct does not adversely affect the differentiation phenotype of T cells through antigen-independent tonic signaling.

### 3.3 Activation and functionality of CAR T cells

To test the functionality of PLAP CAR T cells, we evaluated the expression of CD69 and CD25 activation markers on them upon exposure to PLAP-expressing HeLa cells for 24 hours. PLAP CAR T cells showed a significant increase in the expression of CD69 and CD25 activation markers compared with the control ΔPLAP CAR T cells (Figure 3A).

**Figure 3.**
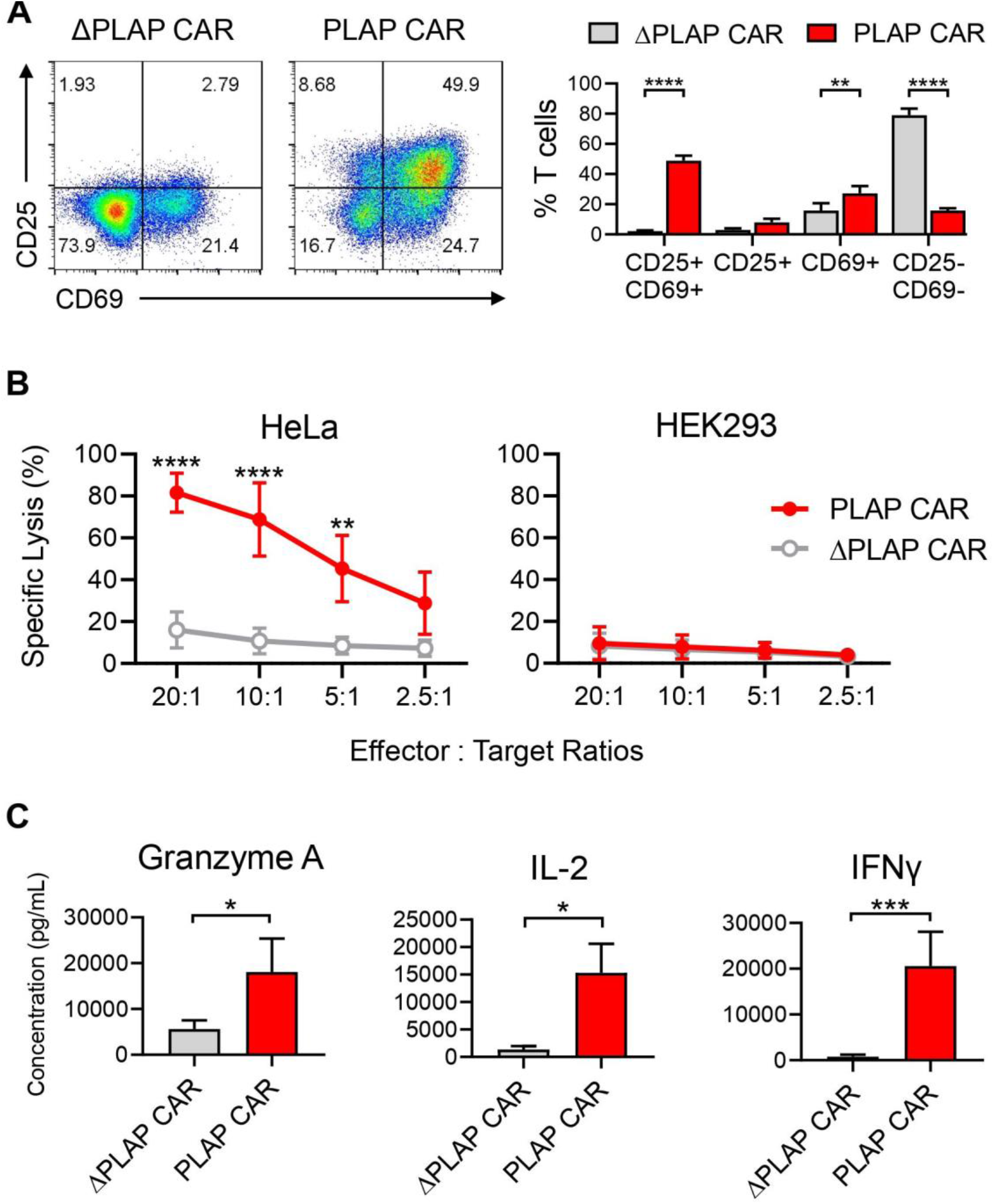
Functional assessment of PLAP CAR T cells. (A) The activation markers such as CD25 and CD69 increase in the PLAP CAR T cells population, but the ΔPLAP CAR T cells population is not double positive. These markers are quantified with anti-CD25 and anti-CD69 by flow cytometry. (B) luciferase-expressing HeLa or HEK293 cells were cocultured with PLAP CAR T cells and ΔPLAP CAR T cells, then by increasing the PLAP CAR T cells, the specific lysis was increased. (C) Secretory factors, including granzyme A, IL-2, and IFN-γ, were measured by ELISA from supernatant and PLAP CAR T cells in comparison to ΔPLAP CAR T cells released higher amounts of anti-tumor molecules. n=3. * P < 0.05; ** P < 0.01; *** P < 0.001; **** P < 0.0001.

Furthermore, by coculturing HeLa cells with different ratios of PLAP CAR T cells and ΔPLAP CAR T cells, we found that by increasing the number of PLAP CAR T cells, the specific lysis also increased (in 20:1 ratio). We observed about 80% specific lysis within two hours. As a control, we used HEK293 cells, which do not express PLAP. We did not observe any significant lysis of HEK293 cells. This demonstrates that the PLAP CAR T cells specifically detect the PLAP antigen expressed on HeLa cells. ΔPLAP has a minor increase in specific lysis against HeLa cells because of the increase in T cells, which shows that non-specific T cells could have a minor effect on tumor regression (Figure 3B).

To evaluate cytokine production by CAR T cells upon activation, we harvested the supernatant 24 hours after coculture of ΔPLAP CAR or PLAP CAR T cells with HeLa cells and measured the granzyme A, IL-2, and IFN-γ secreted into the media. All three cytokines were secreted at higher levels by PLAP CAR T cells compared to ΔPLAP CAR T cells (Figure 3C).

### 3.4 Anti-tumor activity of PLAP CAR T cells

In order to investigate the anti-tumor capacity of PLAP CAR T cells, we co-cultured the PLAP CAR or ΔPLAP CAR T cells with HeLa cells for 9 days. During the course of coculture, PLAP CAR T cells expanded more than 30-fold, reaching around 6.4×10^5^ cells on day 9, significantly higher than the number of ΔPLAP CAR T cells; which was around 1.3×10^5^ on day 9 (Figure 4A and B). More importantly, the number of HeLa cells decreased in the presence of PLAP CAR T cells, while they continued to proliferate in the presence of control ΔPLAP CAR T cells (Figure 4A and 4C). This data demonstrates potent in vitro anti-tumor activity and tumor control ability of PLAP CAR T cells.

**Figure 4.**
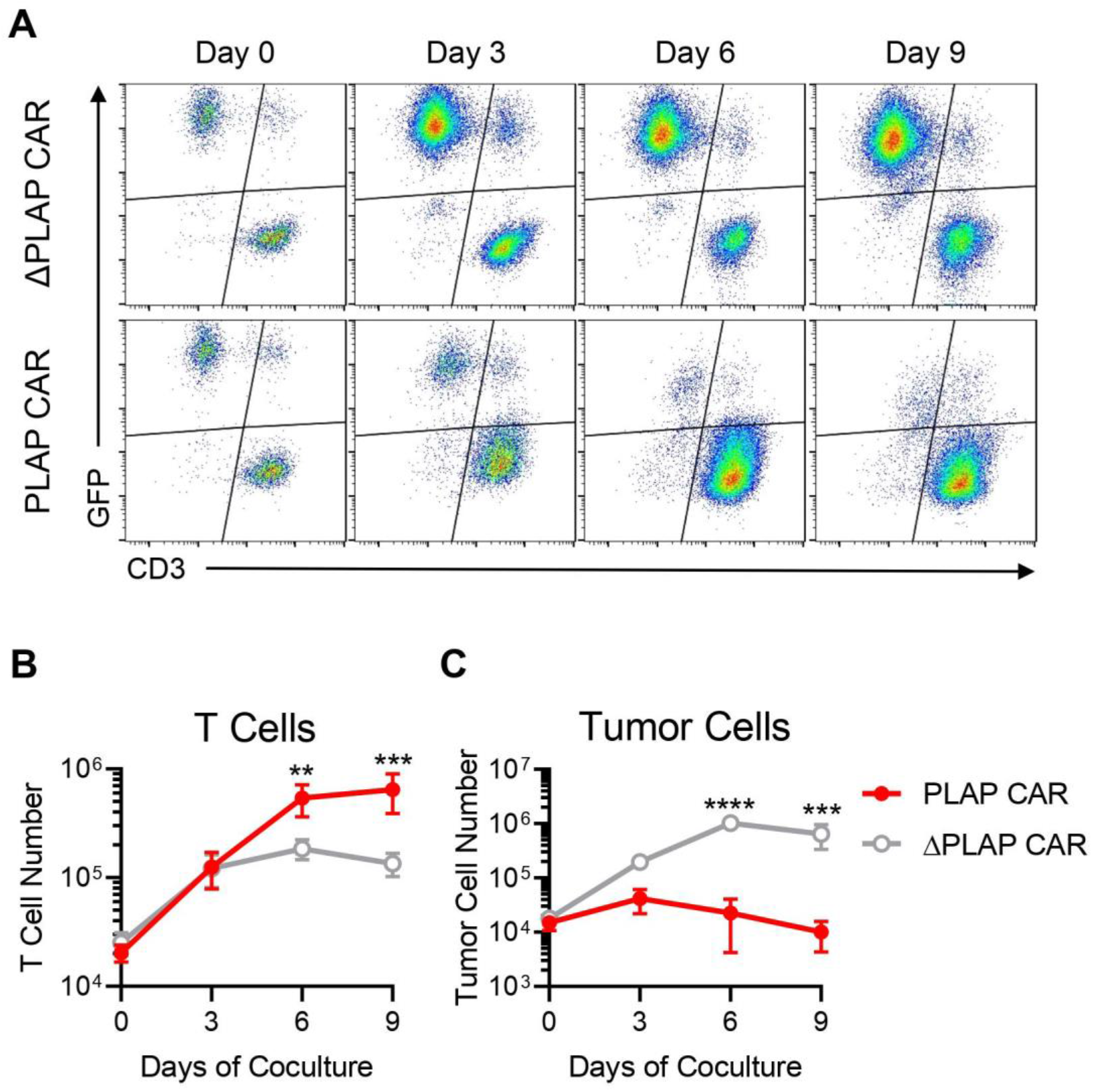
Anti-tumor response of PLAP CAR T cells, in vitro long-term coculture experiment. (A) PLAP CAR T cells and ΔPLAP CAR T cells were cocultured with HeLa cells for 9 days. Cells were collected every 3 days and counted by flowcytometry. PLAP CAR T cells, in comparison with ΔPLAP CAR T cells, demonstrate significant anti-tumor potential, and during 9 days, most of the HeLa cells can be eliminated. (B, C) These graphs show the population of T cells and tumor cells during coculture;(B) PLAP CAR T cells proliferated much more than ΔPLAP CAR T cells, and in (C), the number of HeLa cells which were eliminated in the presence of PLAP CAR T cells was significantly more than when they were cocultured with ΔPLAP CAR T cells. n =3. ** P < 0.01; *** P < 0.001; **** P < 0.0001.

## 4. Discussion

We recently proposed that PLAP can be used as a potent tumor antigen for targeting different solid tumors [18]. High to moderate levels of PLAP expression is only detected in placental tissues, while only a weak PLAP expression has been observed in some endocervical, endometrial, and fallopian tube epithelium samples [11]. This limited PLAP expression in somatic tissues can address the on-target/off-tumor cytotoxicity obstacle in CAR T cell therapy. We designed and constructed a second-generation PLAP-specific CAR and evaluated its functional characteristics by generating PLAP CAR-modified human T cells.

Tonic signaling is a continuous antigen-independent signaling through the CAR molecule resulting in chronic stimulation and early exhaustion of T cells [21, 22]. Signaling through 4-1BB-bearing CARs has been shown to reduce tonic signaling and T cell exhaustion compared with CD28-containing CARs [23]. However, this advantage is vector-dependent. For instance, it is reported that 4-1BB signaling in CD19 CAR T cells, unlike CD28, produces positive feedback on the gammaretroviral long terminal repeat (LTR) promoter through NF-κB activation resulting in CAR T cell apoptosis and inefficient anti-tumor performance [24]. Therefore, we chose to use a CD28 signaling domain in our gammaretroviral PLAP CAR construct. The scFv moiety of the CAR molecule can also contribute to tonic signaling through antigen-independent intermolecular interactions [21, 25]. For instance, in the GD2 CAR T cells, these interactions activate a transcriptional profile that accounts for the expression of inhibitory receptors, which causes exhaustion and apoptosis of GD2 CAR T cells [23]. Thus, it is crucial to confirm that any new scFv-based CAR does not mediate tonic signaling and the associated adverse effects on the T cell function. As CD19 CAR T cells recently showed satisfactory results in hematopoietic malignancies [26, 27], we used them as a standard indicator to compare the PLAP CAR T cells characteristics.

Another factor that has a substantial role in CAR T cells is the type of scFv that should be humanized instead of murine because humanized scFv showed lower immunogenicity compared to murine scFv [28–30], so we used humanized scFv. In a study by Brudno et al., they researched CD19 CAR T cells against B cell lymphoma and found that humanized scFv showed lower neurologic toxicity and cytokine release syndrome (CRS) than murine scFv [31]. Furthermore, one of the drawbacks of murine scFv compared to human scFv is that after injection, they are eliminated by anti-mouse antibodies so, this accounts for lower persistence of CAR T cells [32–35].

T cell subtypes can be distinguished by the expression of the range of CD markers on them, and each of these subtypes exhibits different characteristics like immediate action but low persistency and vice versa [36–38]. In cancer treatment, the population of T cells with less differentiated characteristics like TNL, TSCM, and TCM have more persistency, which is favorable because in the case of cancer relapse, these T cells are required as “living drugs” [39, 40]. Investigation on TCM confirmed they have a better expansion, persistency and anti-tumor effect in the long-term, *in vivo* [41–43]. Moreover, according to figure 2E, the population of TCM is higher than other subtypes of CAR T cells. Another indicator that shows PLAP CAR T cells are less differentiated than other groups is the high expression of CD27 and CD28 on PLAP CAR T cells (Figure2D), which can stimulate them against cancer cells [44, 45].

During the activation of CAR T cells, they release anti-tumor molecules, including interleukins, perforin, granzyme [46, 47] and other markers [48]. By harnessing these characteristics, we showed that during coculture with target cells PLAP CAR T cells, secrete IFN-γ, granzyme A, and IL-2 and express CD25 and CD69; along with specific lysis of PLAP^+^ cells which altogether confirmed the functionality of PLAP CAR T cells (Figure3). Besides, Li et al. observed specific lysis and release of IFN-γ; when they co-cultured PLAP CAR T cells with colon cancer cells (PLAP^+^ cells) [49].

The serial killing of PLAP CAR T cells was confirmed by coculturing with target cells in the long term and comparing their killing capacity with ΔPLAP CAR T cells. Although establishing the potential of PLAP CAR T cells in tumor regression needs to be evaluated in animal models, but another experiment published during our research showed that PLAP CAR T cells in mice model with colon cancer resulted in tumor regression [49].

## 5. Conclusions

Here we demonstrate the importance of constructing CAR T cells against novel antigens that can be a potential asset in solid tumor regression. PLAP CAR T cells can be a prospective weapon in targeted cancer immunotherapy, which needs more investigation on other PLAP^+^ cancer cells to find their own way in clinical trials.

## Author Contributions

Conceptualization, S.P, and M.B.; methodology, V.Y, S.P, A.S, and M.B.; data analysis, V.Y, A.S, Z.M, and M.B; writing—original draft preparation, V.Y, and M.B.; writing—review and editing, V.Y, S.P, Z.M, and M.B; visualization, S.P, and M.B.; supervision and funding acquisition, B.M, and M.B.

## Funding

This research was supported by funds from (a) the Royan institute and (b) Science and Technology Commission of Shanghai Municipality (20ZR1426500 to B.M)

## Institutional Review Board Statement

The study was conducted in accordance with the Declaration of Helsinki, and approved by the Research Ethics Commitee of Royan institute (reference number IR.ACECR.ROYAN.REC.1397.112 approval).

## Informed Consent Statement

Informed consent was obtained from all blood donors involved in the study.

## Data Availability Statement

The data presented in this study are available on request from the corresponding author.

## Acknowledgments

We thank all staff and technicians at Royan Institute for Stem Cell Biology and Technology for their technical assistance.

## Conflicts of Interest

The authors declare no conflict of interest.

## Notes

### Competing Interest Statement

The authors have declared no competing interest.

